# Hydrogel-coating improves the in-vivo stability of electrochemical aptamer-based biosensors

**DOI:** 10.1101/2020.11.15.383992

**Authors:** Shaoguang Li, Jun Dai, Man Zhu, Netzahualcóyotl Arroyo-Currás, Hongxing Li, Yuanyuan Wang, Quan Wang, Xiaoding Lou, Tod E. Kippin, Shixuan Wang, Kevin W. Plaxco, Hui Li, Fan Xia

## Abstract

The ability to track the levels of specific molecules, such as drugs, metabolites, and biomarkers, in the living body, in real time and for long durations would improve our understanding of health and our ability to diagnose, treat and monitor disease. To this end, we are developing electrochemical aptamer-based (E-AB) biosensors, a general platform supporting high-frequency, real-time molecular measurements in the living body. Here we report that the addition of an agarose hydrogel protective layer to E-AB sensors significantly improves their baseline stability when deployed in the complex, highly time-varying environments found in vivo. The improved stability is sufficient that these hydrogel-protected sensors achieved good baseline stability when deployed in situ in the veins, muscles, bladder, or tumors of living rats without the use of the drift correction approaches traditionally required in such placements. Finally, this improved stability is achieved without any significant, associated “costs” in terms of detection limits, response times, or biocompatibility.

## Introduction

The ability to track the levels of specific molecules, such as drugs, metabolites, or biomarkers continuously and in real time in the living body would vastly improve our knowledge of physiology, pharmacokinetics, and toxicology and would pave the way for truly high-precision personalized medicine. Such a technology, for example, would improve our understanding of many time-dependent physiological events, including the distribution phases of therapeutic drugs, the pulsatile release of hormones, and the hemostatic control of key metabolites.^[1–3]^ In the clinic such a technology could likewise provide the high-precision, patient-specific pharmacokinetics required to provide high precision dosing, and even support real-time, feedback-controlled drug delivery of unprecedented accuracy.^[4–5]^ The development of such a technology, however, faces significant hurdles.^[6–7]^ Specifically, such a technology must: (1) achieve clinically-relevant sensitivity and specificity; (2) must be reversible, so that it can follow rising and falling concentrations; (3) it must operate continuously, or at least at a frequency that is high relative to physiological timescales (it thus cannot rely on batch processing, such as separations or the addition of exogenous reagents); and (4) it must remain stable in the complex, fluctuating environments found within the body. Faced with these hurdles, continuous, in-vivo sensing had, until recently only been reported for a short list of metabolites, physiological molecules (e.g., glucose,^[8]^ lactate,^[9]^ and blood oxygen^[10]^) and neurotransmitters (e.g., dopamine, acetylcholine).^[11–12]^ Moreover, the sensors for each of these targets are critically reliant on the chemical or enzymatic reactivity of their targets, and thus they are not generalizable to the detection of other, arbitrary targets.

Against this background, we^[6, 12–14]^, followed by others^[15–17]^, have developed electrochemical aptamer-based (E-AB) biosensors, the first platform technology supporting high-frequency, in-vivo molecular measurement that does not rely on the intrinsic chemical or enzymatic reactivity of its targets. To achieve this, E-AB sensors employ a target binding-induced conformational change to generate an electrochemical signal (**Fig. 1a**). Specifically, E-AB sensors are comprised of redox-reporter-modified DNA or RNA aptamer, a class of functional oligonucleotides that binds a specific analyte and can be artificially selected via high-throughput, in-vitro methodologies, that are covalently attached to an interrogating electrode. The binding of an analyte to these recognition elements alters the efficiency with which the redox reporters transfers electrons to or from the electrode, producing an easily measurable signal change when the sensor is interrogated electrochemically.^[12]^ Because this signaling mechanism recapitulates the conformation-linked signaling typically employed by naturally occurring chemo-perception systems, E-AB sensors are selective enough to deploy directly in complex sample matrices.^[18–19]^

**Fig. 1.**
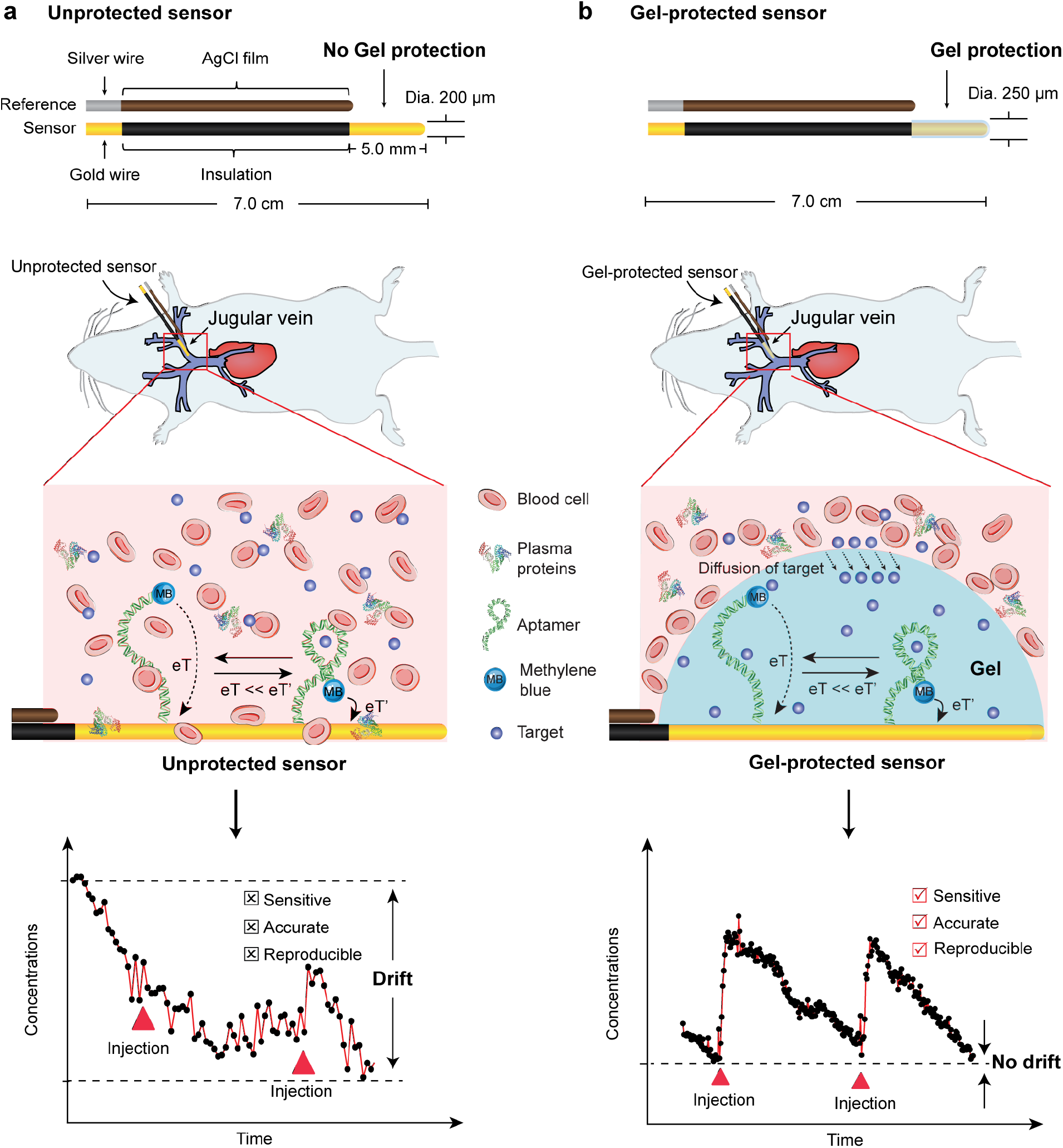
Here we have demonstrated the ability of a protective hydrogel coating layer to improve the baseline stability of electrochemical aptamer-based (E-AB) biosensors when they are deployed in situ in the living body. The signaling mechanism of E-AB sensors, which recapitulates the conformation-linked signaling employed by naturally occurring chemo-perception systems, renders them selective enough to be deployed in complex matrix, such as blood serum.^[18–19]^ (**a**) However, they suffer from significant signaling baseline drift when they are deployed directly in whole blood for continuous, real-time measurements both in vitro and in vivo without any drift corrections (bottom panels shown are real data collected in vivo). Historically we have corrected this drift using algorithms such as kinetic differential measurements or chronoamperometry.^[23–24]^ (**b**) Here, in contrast, we demonstrate the good baseline stability of uncorrected, gel-protected sensors (bottom panels shown are real data collected in vivo).

While E-AB sensors are reasonably stable in blood and serum in vitro, they exhibit significant drift when deployed directly in the living body^[20–22]^, which presumably arises due to degradation of the target-recognizing aptamer and the non-specific adsorption of cells and other blood components to the sensor surface. To circumvent this, we have historically employed drift correction methods, such as kinetic differential measurements (KDM)^[23]^, dual-reporter approach^[12]^ or chronoamperometry.^[24]^ The former two employs a secondary signal from the sensor to correct for the drift, while the latter monitors the electron transfer time constant to quantify the target, an observation that is independent of the number of aptamer probes and thus rather insensitive to sensor degradation. Using these approaches, we have achieved good measurement accuracy and returns to baseline for in-vivo runs up to several hours. Ultimately, however, these approaches fail when the sensor degradation becomes so great that its signal-to-noise ratios become unacceptably low. Thus motivated, here we explore a complementary technology: the use of a hydrogel matrix to protect E-AB sensors from non-specific adsorptions and sensor degradations by decreasing the access of high molecular weight, biological components (e.g., proteins, blood cells) to the sensor surface (**Fig. 1b**), reducing their ability to degrade sensor performance.

## Results

### Gel-coating improves E-AB sensor performance

The utility of gel coating to solve the baseline drift of E-AB sensors relies on the differential kinetics of molecular diffusion through a dense hydrogel. To characterize this diffusion, we first used both modeling and experimental approaches to estimate the size-dependence of molecular diffusion kinetics through a representative hydrogel. Specifically, using Amsden’s theoretical model, ^[25]^, we found that the time required for diffusion through a certain length of hydrogel is greatly dependent on both molecular size (here, we employed a variety of proteins in whole blood) and gel porosity (i.e., gel wt% concentrations, **Fig. 2a** and **2b**, Supplementary Fig. 1). Consistent with this, when we incubated a 5 mg/mL solution of fibrinogen (~25 nm in diameter) and 5 mg/mL solution of BSA (~3 nm in diameter) on top of a gel for 24 h and monitored the two proteins’ diffusion into the gel mass, we found that, at a depth of ~25 μm (the thickness of the gel on our protected sensors), the concentration of the former only reached 10% of the applied concentration (**Fig. 2c and 2d**). In contrast, the latter, lower-molecular weight protein was homogeneously distributed throughout the entire ~ 400 μm thickness of the gel slice.^[26–27]^ Not surprisingly, the presence of a hydrogel also prevents the agglomeration of blood cells onto a sensor surface (**Fig. 2e**), as these are far too large to penetrate the gel network.

**Fig. 2.**
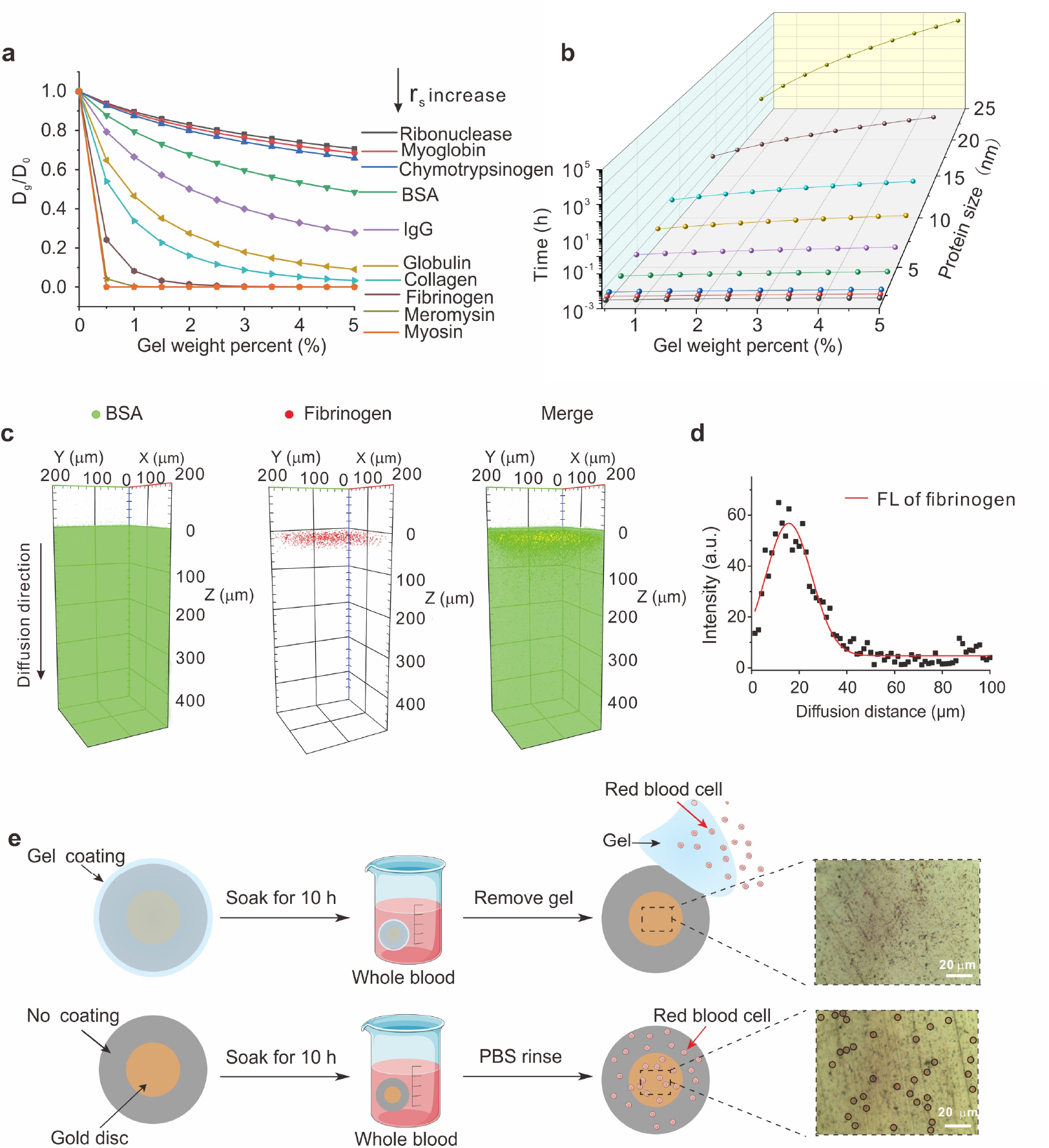
Hydrogel greatly reduces the diffusion of high-molecular weight proteins and effectively eliminates the diffusion of cells. (a) Using Amsden’s theoretical model, we modeled the extent to which the presence of a hydrogel reduces the diffusion constant of various proteins (D_g_/D_0_ reflects the diffusion constant in the gel relative to that seen in the free solution). As expected, the gel significantly reduces the diffusion of high-molecular weight proteins, while have less of an effect on lower molecular weight proteins. (b) The time duration required for blood components to diffuse through a specific depth (here, ~25 μm) is likewise greatly dependent on their molecular weight and the density of the gel. (c-d) Using confocal microscopy we have monitored the diffusion of BSA (~3 nm in diameter) and fibrinogen (~25 nm) through a gel slice fabricated from 3 wt% gel solution over the course of 24 h. Under these circumstances the concentration of fibrinogen at a depth of 25 μm reaches only ~10% of the applied concentration. In contrast, after the same 24 h incubation the concentration of BSA seen in the gel is homogeneously distributed across its entire 400 μm thickness of the gel. (e) Not surprisingly, while red blood cells adhere to an unprotected electrode (i.e., no gel coating) none are seen on a gel-protected electrode (The red blood cells on the unprotected sensors are outlined with black circles).

### Baseline stability in-vitro

Motivated by the ability of a hydrogel to greatly reduce the diffusion of blood cells and higher molecular weight proteins, we next investigated their ability to protect E-AB sensors in vitro in (a) undiluted whole blood and (b) excised, solid tissue, using a sensor against the antibiotic kanamycin^[14]^ as our test bed. Using a standard dip-coating protocol (Supplementary Fig. 2), we first fabricated a gel-protected E-AB sensors and then interrogated these sensors in vitro both in buffer and whole blood using a three-electrode system (Fig. 3a). As expected, we observed similar stability when we challenge unprotected and gel-protected versions of this sensor in phosphate buffered saline (Supplementary Fig. 3). In contrast, when the two are challenged in vitro in undiluted whole blood, the performance of the gel-protected sensor is notably improved over that of unprotected sensors. For example, over 10 h the peak currents of unprotected sensors fall by 70% in undiluted whole blood and over 10 h under these conditions, those of gel-protected sensors fall by less than 5% (**Fig. 3b, c, d**). Perhaps not coincidentally, after interrogating for 10 h in whole blood, we observed that, while unprotected sensor was covered with biocomponents, gel-protected sensor exhibited no adsorptions to such components (Supplementary Fig. 4). The improved stability of gel-protected sensors likewise holds for sensors inserted into excised solid tissue (Fig. 3e) and for sensors employing aptamers against other than kanamycin (Supplementary Fig. 5).

**Fig. 3.**
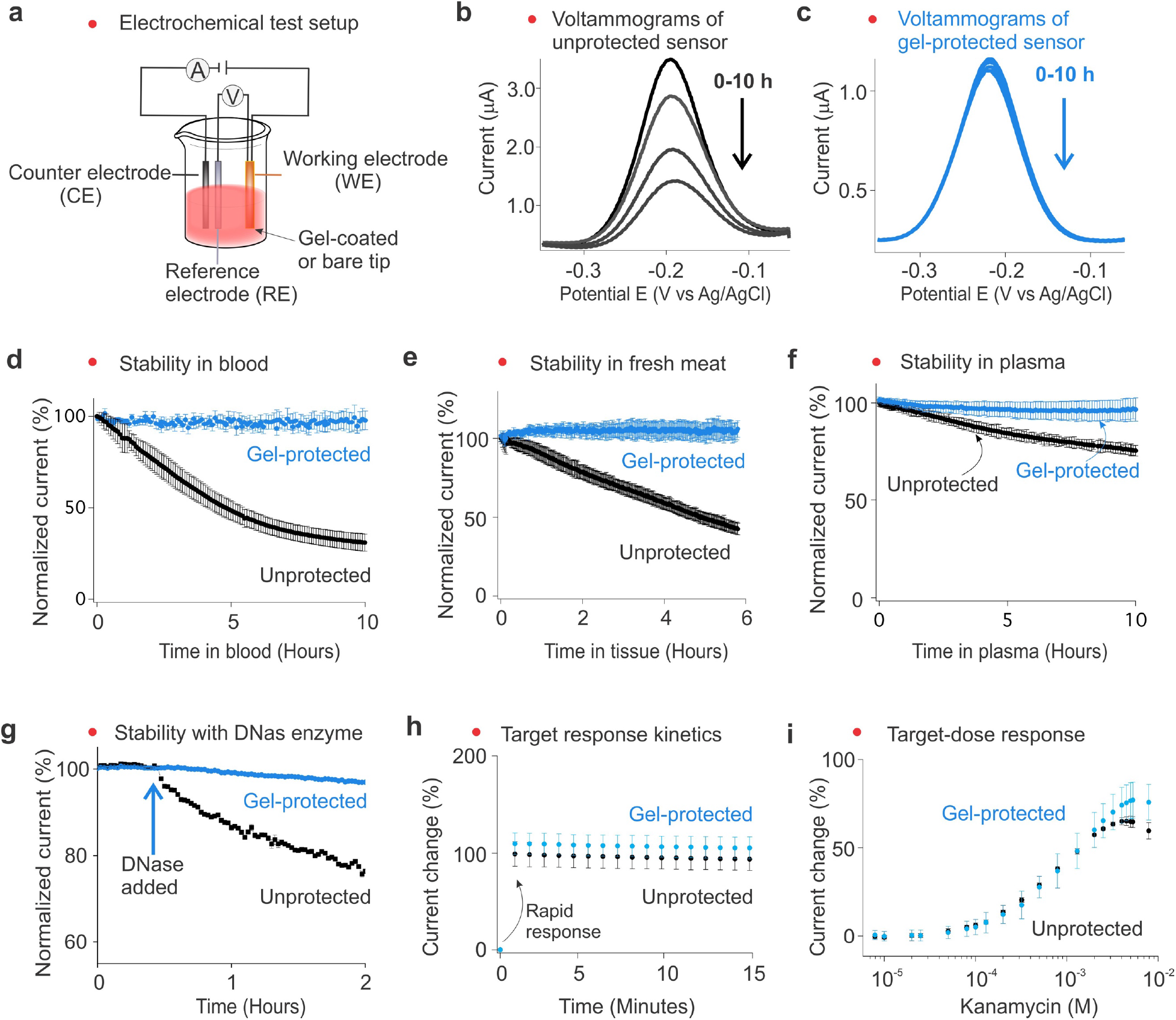
Gel-protected sensors achieve excellent stability while maintaining rapid response time and low limits of detection. (a) Here we employed a three-electrode system to interrogate our sensor, including a gold working electrode that serves as the sensor, a Ag/AgCl reference electrode, and a platinum counter electrode. The gold working electrode was used as is or protected by gel layer, i.e., unprotected or gel-protected sensors. (b) When deployed in whole blood, unprotected sensors exhibited a significant signal loss shown as the current obtained via square wave voltammetry decaying dramatically. (c) Under the same conditions, the signal obtained from gel-protected sensors is quite stable. (d) For example, while unprotected sensors exhibit 80% signal loss over 10 h in whole blood, gel-protected sensors exhibit no signal loss under the same conditions. After interrogating for 10 h in whole blood, while unprotected sensor was fully covered with biocomponents, gel-protected sensor exhibited no adsorptions to these components (Supplementary Fig. 4). Gel-protected sensors likewise remain far more stable (e) when inserted into a tissue sample (here fresh pork) and (f) in plasma. (g) When deployed in PBS buffer in the presence of a 5 mM solution of DNases-I, unprotected sensors exhibit 20% signal loss over 1.5 h, while for gel-protected sensors the loss is less than 5%. The (h) response time and (i) limits of detection of gel-protected sensors remain quite similar to those of unprotected sensors.

To determine which blood components contribute most significantly to the observed drift we next challenged gel-protected and unprotected sensors for 10 h in either plasma, which is the liquid fraction of whole blood (**Fig. 3f**), or formed elements (Supplementary Fig. 6), which is the cellular fraction. When incubated for 10 h, gel-protected sensors exhibited excellent stability under both conditions. In contrast, unprotected sensors lost 80% of their signal in formed elements and 20% in plasma, suggesting that the agglomeration of blood’s cellular components on the sensor or the reaction of their components with the DNA aptamer (e.g., DNAses liberated from ruptured cells, **Fig. 3g**) are a larger source of baseline drift than that associated with the proteins in plasma.

The drift protection provided by the gel coating comes at relatively little cost in sensor equilibration time, detection limits. For example, while the presence of the protecting gel slows sensor response times, this effect is small relative to the timescales of the physiological processes E-AB sensors have been used to investigate. Specifically, a kanamycin-detecting, gel-protected sensor reaches 90% of its maximum signal change within less than one minute (**Fig. 3h**), a timescale far faster than the tens of minutes elimination rate of this drug.^[14]^ The detection limit of a gel-protected sensor is likewise effectively indistinguishable from that of the equivalent, unprotected sensor (**Fig. 3i**). Consistent with this, the electron transfer kinetics of gel-protected sensors are closely comparable to those observed for unprotected sensors (Supplementary Fig. 7), suggesting their gain and detection limits should be similar.

### Continuous, real-time, in vivo molecular measurements

Motivated by the improved stability of gel-protected sensors exhibit in vitro, we next tested them in a variety of in vivo scenarios. As the first of these tests we emplaced a kanamycin-detecting sensor in the external jugular veins of an anesthetized Sprague-Dawley rats, and injected the drug into the opposite external jugular vein. Under these conditions, an unprotected (non-drift-corrected) sensor exhibits ~40% loss in signal over the course of 2 h (**Fig. 4a and 4b**). Gel protection reduced this loss to less than 5% over the same period. Following two sequential intravenous boluses of kanamycin, the gel-protected sensor recorded consecutive concentration spikes corresponding to each bolus, with maximum kanamycin concentrations (C_max_) of ~200 μM and the effective clearance of 90% of the drug from the circulatory system within 50 min (**Fig. 4c**), values that are fully consistent with prior studies of the pharmacokinetics of this drug.^[6]^ Repeating these measurements with the gel-protected sensor in multiple rats we observed reproducibility consistent with the known pharmacokinetic variability of the aminoglycosides^[28]^ (Supplementary Fig. 8-9). Finally, we used gel-protected sensors to follow monotonically increasing intravenous kanamycin doses spanning the 10-30 mg/kg therapeutic ranges used in humans^[29]^ (**Fig. 4d**). The sensor responded rapidly to each injection, and measured peak drug concentrations in good accordance with the relevant delivered dose (**Fig. 4e**).

**Fig. 4.**
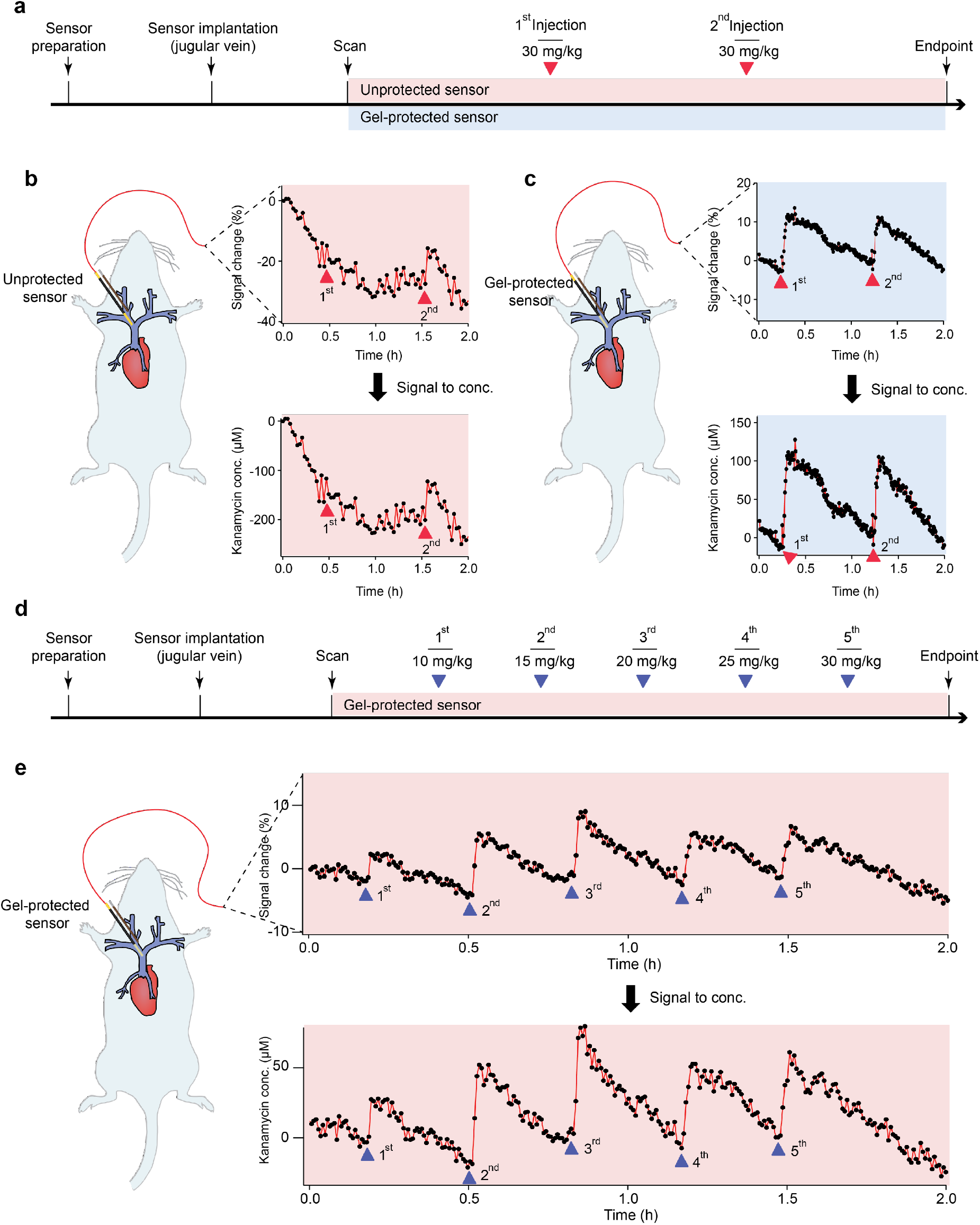
Gel-protected E-AB sensors support continuous, real-time molecular measurements in situ in the living body without the use of the drift-correction algorithms previously used in such deployments.^[23]^ (a) To show this, we first emplaced kanamycin-detecting sensors in the external jugular veins of anesthetized Sprague-Dawley rats, which we then challenged via intravenous injection of the drug into the opposite external jugular vein. (b) Unprotected sensors exhibited significant drift over the course of these few-hour experiments, rendering them incapable of determining the target concentrations without employing drift-correction methods. (c) In contrast, gel-protected sensors are much more stable, achieving precise, continuous molecular measurement. (d) We then followed monotonically increasing intravenous doses of kanamycin spanning the 10-30 mg/kg therapeutic ranges used in humans. (e) The sensor responded rapidly to each injection, measuring maximum concentrations between 34 and 100 μM in good accordance with the delivered dose.

Gel-protected sensors achieve clinically relevant accuracy in vivo without employing the drift correction mechanism we have previously employed. To see this, we performed simultaneous in-vivo and ex-vivo measurements using, respectively, a gel-protected E-AB sensor and blood draws followed by high-performance liquid chromatography (**Fig. 5**). Specifically, we implanted one kanamycin-detecting, gel-protected sensor in the jugular vein of a living rat and performed the real-time, continuous E-AB measurements while drawing blood samples at 20 to 30 min intervals from the opposite jugular vein. Performing HPLC-ELSD measurements immediately after in-vivo test (to maintain the freshness of the sample) using a standard protocol for blood pre-treatment (detailed protocols see supplementary materials, Supplementary Figs. 10-11), the ex-vivo measured concentrations for each sample were in close accordance to the values obtained from E-AB sensors (**Fig. 5b, c**). For example, measurements taken by the two approaches during the elimination phase of the drug’s pharmacokinetics were within 30% of one another, a level of accuracy similar to that of commercial glucose sensors.^[30–31]^

**Fig. 5.**
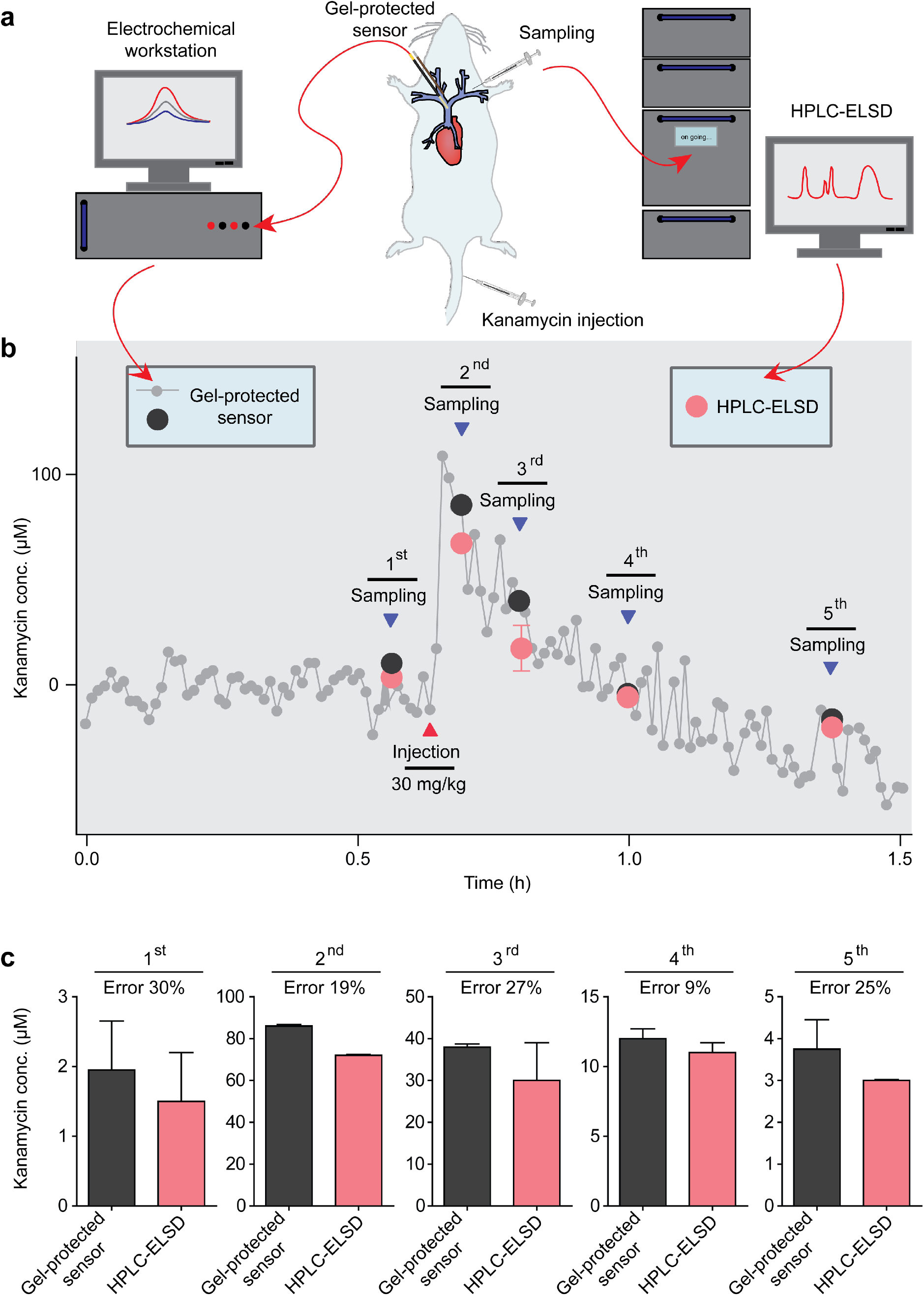
Gel-protected E-AB sensors achieved clinically relevant accuracy without employing drift-correction approaches.^[23–24]^ (**a**) and (**b**) To see this, we performed parallel pharmacokinetics studies using ex-vivo analysis via high-performance liquid chromatography as our gold standard. Specifically, used a gel-protected E-AB sensor in the jugular vein of a live rat to perform real-time measurements of kanamycin, while simultaneously collecting blood samples at an interval of 20 to 30 minutes for subsequent bench-top analysis. (**c**) The results of the two approaches are in good accordance throughout the experiment. The precision of E-AB sensors as defined by the standard deviation observed during the pre-challenge baseline is on the order of ~30 μM. The error bars on the E-AB data are derived from the error in the calibration curves obtained via in-vitro titrations. The error bars on HPLC data were derived from replicate measurements performed on each sample. In both cases the error bars reflect 95% confidence intervals.

### Simultaneous, multi-compartment measurements

Gel-protected E-AB sensors also achieve good baseline stability when placed in other bodily compartments, suggesting that they can be used, for example, to monitor time-varying molecular concentrations throughout the body^[32–33]^ To see this, we first implanted three E-AB sensors in a single rat: one in a jugular vein, a second in a leg muscle, and a third in the bladder (**Fig. 6a**). Upon intravenous challenge with 40 mg/kg kanamycin, we again observed a rapid increase in plasma drug level peaking at ~200 μM. As expected, the sensors placed in the tissue and bladder also measured rising drug levels, albeit with a delay arising due to the slow transport of the drug into the tissues and the slow excretion of the drug via the kidneys (**Fig. 6b**). Following on this, we also implanted a sensor against the chemotherapeutic doxorubicin into a solid tumor (**Fig. 6c**, Supplementary materials, **video 1**). Once again, when we implanted a doxorubicin-detecting gel-protected sensor in BALB/C nude mice, we achieve a micromolar precision upon intravenous challenge with the drug (**Fig. 6d**) at therapeutically relevant concentrations.

**Fig. 6.**
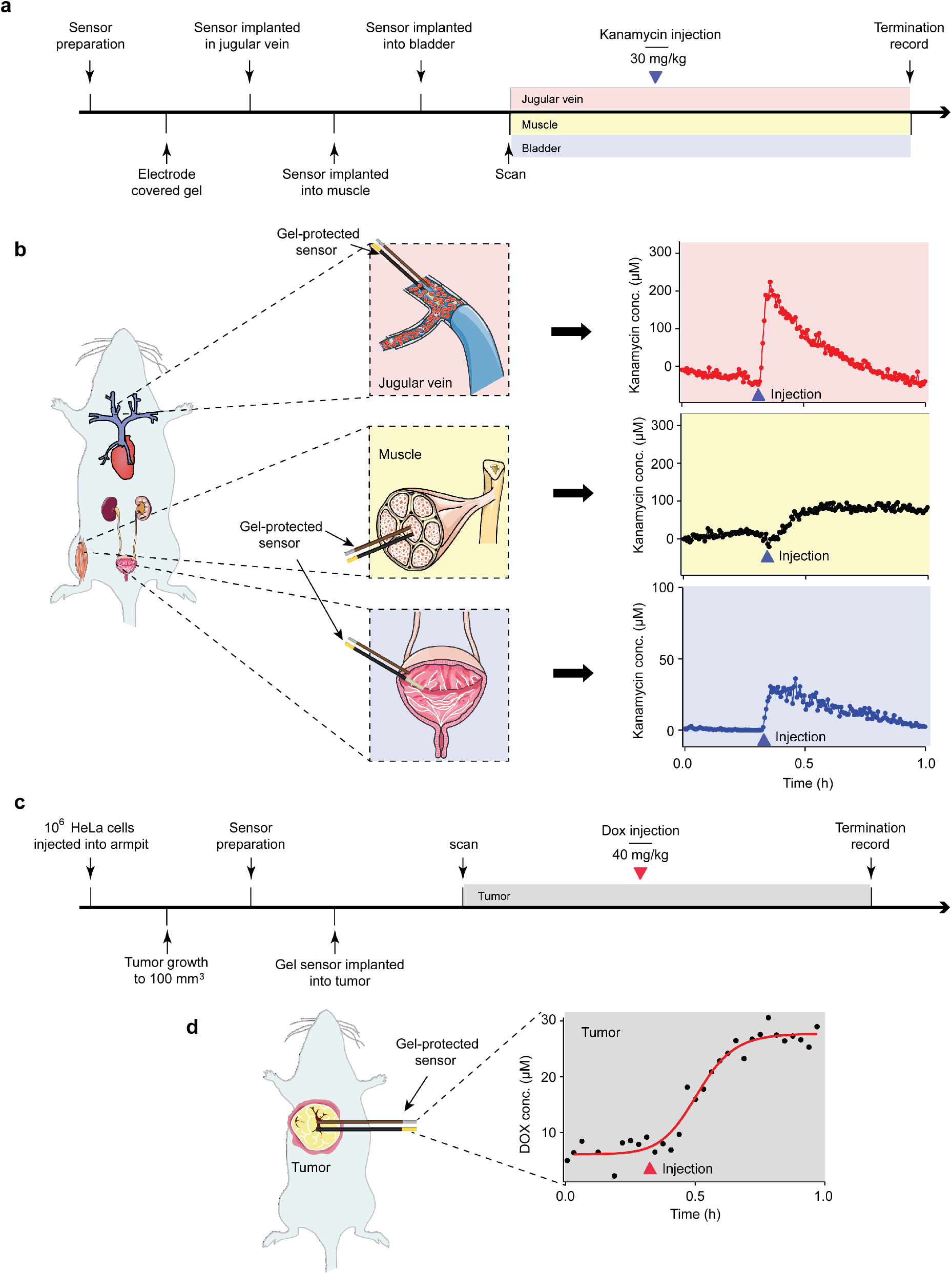
Gel-protected E-AB sensors exhibit good baseline stability when placed in a variety of bodily compartments. (a) To show this we simultaneously implanted kanamycin-detecting sensors in the jugular vein, leg muscle, and bladder of a single rat. (b) Upon intravenous challenge with 40 mg/kg kanamycin we once again observed an immediate rise of drug in the vein, peaking at ~200 μM. The sensors in the muscle and bladder exhibited slower drug level rise, indicating a slower uptake into the solid tissues and slower elimination of the drug from the blood. (c) We have also explored the use of E-AB sensors to perform continuous, real-time molecular tracking in situ in a tumor. (d) Specifically, a gel-protected, doxorubicin-detecting sensor placed inside of a xenograph tumor achieved micromolar precision upon an intravenous challenge of the drug at 40 mg/kg, a dose that is therapeutically relevant.

### Biocompatibility

We also performed preliminary exploration of the biocompatibility of both unprotected and gel-protected E-AB sensors, a critical issue for the development of long-duration implantable devices.^[7, 34–37]^ To do so we employed five groups of rats (15 rats per group): those implanted with (1) gel-protected and (2) unprotected sensors in the tail vein; those implanted with (3) gel-protected and (4) unprotected sensors in the muscle of the left hind limb; and (5) a control group that was not implanted with any sensors, but was subjected to the same surgical procedures as a combination of both vein and muscle groups. In each case, the sensors remained in the animal for one week. During this time observed no significant difference among the five groups in terms of body weight, food/water consumption, or behavior (**Fig. 7a-c**). A week post-implantation we performed blood examination and morphology studies. Upon visual inspection, we observed what appeared to be complete recovery of the implantation site and no inflammation of the skin (Supplementary Figs. 12-17). Consistent with this, we did not observe any significant differences in blood vessel morphology, and only slight differences in muscle tissue between implanted sensors (either protected or unprotected) and controls (**Fig. 7d**). Likewise, we did not detect any hematological changes in terms of white blood cells, red blood cells, hemoglobin, and platelets counts (**Fig. 7f**. To test for the presence of thrombosis in intravenous placements,^[38]^ we collected the vessels from the rats one week after tail-vein implantation. In neither case did we observe thrombosis for either gel-protected or unprotected implantations (**Fig. 7g**, Supplementary Fig. 18). Neither did we detect any significant differences in endothelin, D-dimer and thrombomodulin between the five groups (**Fig. 7h**), further suggesting that sensor implantation does not lead to thrombosis. Finally, to evaluate the immune response provoked by our sensors (which can be an obstacle to practical implantable devices^[39–40]^), we monitored the inflammatory biomarkers C-reactive protein, tumor necrosis factor α, interleukin 12 and interleukin 6, observing no statistical difference (*p* >0.05) between any of the five groups (**Fig. 7i**).

**Fig. 7.**
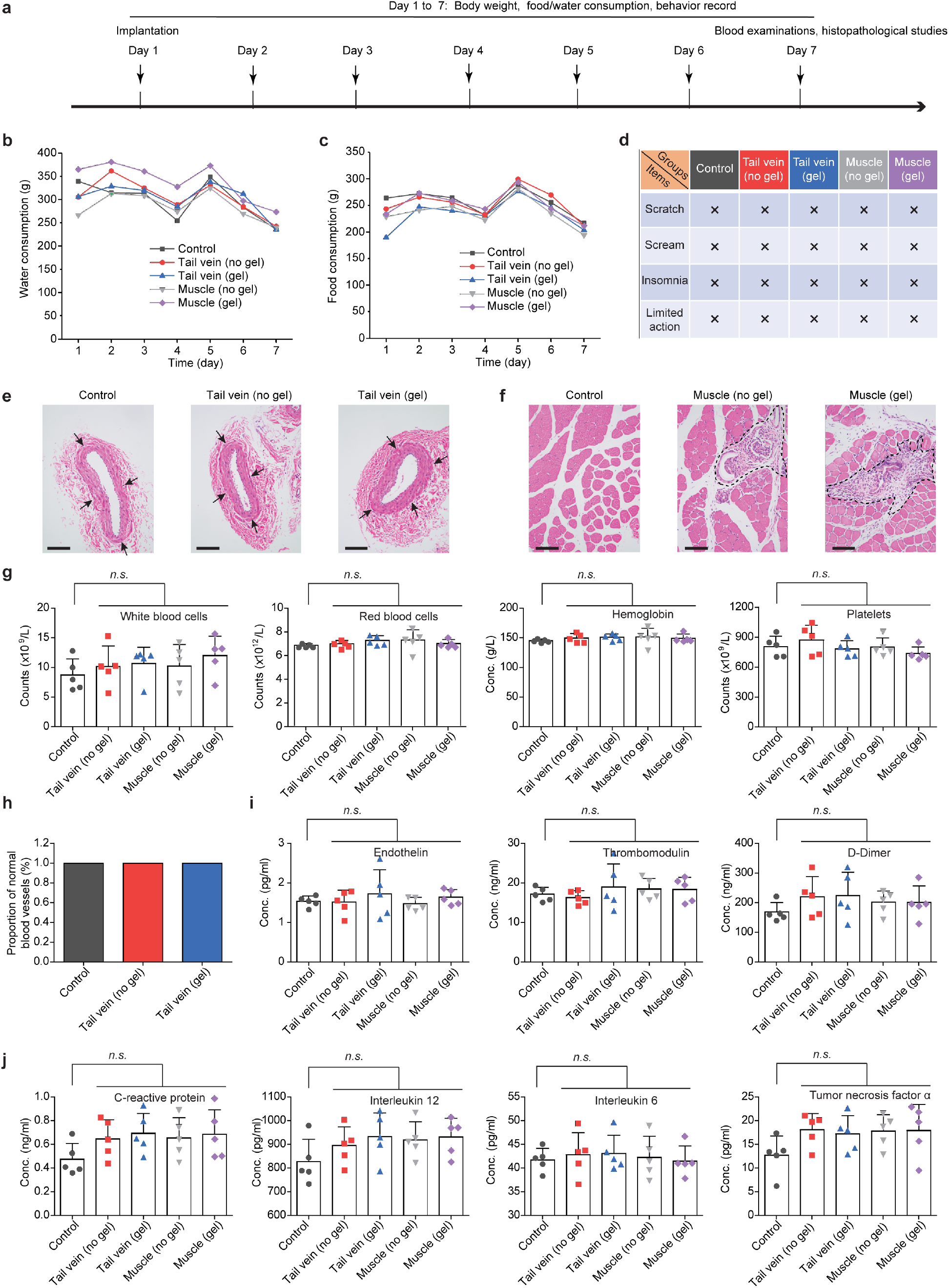
Both gel-protected and unprotected sensors exhibit good biocompatibility. To determine this we employed five groups of 15 rats each: (1) gel-protected and (2) unprotected sensors (denoted here as “no gel”) implanted in the tail vein; (3) gel-protected and (4) unprotected sensors implanted in the muscle of the left hind limb; and (5) a control group that was not implanted with any sensors, but was subjected to the same surgical procedures (both vein and muscle) as the former groups (Supplementary Figs.12-17). (We observed no significant differences in (a) water consumption, (b) food consumption or (c) animal behavior between implanted and control animals. (d) Likewise, we observed no significant difference in the morphology of the tail vein after one-week implantations. Here, the black arrows denote the tail vein. (e) Implantations of both protected and unprotected in the muscle cause mild inflammation, as indicated by black dotted lines. (f) We observe no significant differences in blood counts between any of these five groups. Likewise, we observed (g) no thrombosis, and (h) no significant change in the coagulation markers endothelin, D-dimer, or thrombomodulin. (i) Finally, we observed no sigificant differences in the plasma levels of the inflammatory factors C-reactive protein, interleukin 12, interleukin 6 and tumor necrosis factor α.

## Conclusions

Here we demonstrate that the application of an argarose hydrogel coating significantly reduces drift when E-AB sensors are deployed both in vitro in undiluted whole blood and in situ in the veins, the bladder, solid healthy tissue, or solid neoplastic tissues of live rats. We believe this improved stability is due to the dependent differential kinetics of molecule diffusion across the gel, which allows low-molecular weight target molecules to diffuse rapidly to the sensor surface while largely blocking the approach of high-molecular weight components, minimizing non-specific adsorption to the sensor and enzymatic degradation of its aptamer. Moreover, gel-protected sensors achieve these improvements without significant reductions in time resolution, limits of detection, precision, or accuracy. Finally, we have demonstrated that gel-protected sensors retain the same biocompatibility as unprotected E-AB sensors when implanted in blood vessels or muscle tissue, as neither causes any detectable impact on animal behavior or blood properties, and only minimal changes in tissue morphology.

## Acknowledgements

This work was supported by the National Natural Science Foundation of China (21525523, 21804121, 21874121, 21801231), the Fundamental Research Funds (CUG170665, CUG170668) for the Central Universities, China University of Geosciences (Wuhan) and the US National Institutes of Health (EB022015).

## Author contributions

Dr. S. L. and Dr. H. L. conceived of the project. Dr. S. L., Dr. H. L., Dr. K.W. P., Dr. J. D., Dr. X. L. and Dr. F. X. designed the experiments. M. Z., H. L fabricated the sensors and tested the sensors, conducted the HPLC measurements. Dr. S. L., Dr. J. D., Dr. H. L., Q. W. and Dr. N. A.-C. directed the animal studies, performed the controller simulations and analyzed the data. Dr. H. L., Dr. K.W. P., Dr. S. L. and Dr. J. D. wrote and edited the manuscript. All authors discussed the results and commented on the manuscript.

## Additional information

Supplementary information is available for this paper.

