## Supplementary Material for "Hydrogel-coating improves the in-vivo stability of electrochemical aptamer-based biosensors"

**Table of contents**

Materials.......... S4

Gold electrode preparation and cleaning.......... S4

The fabrication of unprotected and gel-protected sensors.......... S5

Diffusion model.......... S5

Gel electrophoresis.......... S6

Confocal microscopy studies.......... S6

Optical microscopy studies.......... S7

Electrochemical measurements.......... S7

Target-binding equilibration measurements.......... S7

Target titration measurements.......... S8

In vivo experiments.......... S8

Anesthesia.......... S8

Venous sensor implantation.......... S8

Bladder sensor implantation.......... S8

Muscle sensor implantation.......... S9

Tumor sensor implantation.......... S9

HPLC-ELSD measurements.......... S9

Biocompatibility assessment.......... S10

Histopathological examination.......... S10

Blood test.......... S10

ELISA assays.......... S10

Supplementary Fig. 1.......... S11

Supplementary Fig. 2.......... S11

Supplementary Fig. 3.......... S12

Supplementary Fig. 4.......... S12

Supplementary Fig. 5.......... S13

Supplementary Fig. 6.......... S13

Supplementary Fig. 7.......... S14

Supplementary Fig. 8.......... S15

Supplementary Fig. 9.......... S16

Supplementary Fig. 10........................................................................................................ S17
Supplementary Fig. 11........................................................................................................ S17
Supplementary Fig. 12........................................................................................................ S18
Supplementary Fig. 13........................................................................................................ S18
Supplementary Fig. 14........................................................................................................ S19
Supplementary Fig. 15........................................................................................................ S19
Supplementary Fig. 16........................................................................................................ S20
Supplementary Fig. 17........................................................................................................ S20
Supplementary Fig. 18........................................................................................................ S21
Reference.............................................................................................................................. S21

### Materials

Biowest regular agarose G-10 was purchased from Solarbio Science & Tecnology Co., Ltd. (Beijing, China). Trifluoroacetic acid (TFA) was purchased from Beijing InnoChem Science &Technology Co., ltd. (Beijing, China). Bovine serum was purchased from Shanghai Yuanye Bio-Technology Co., Ltd. (Shanghai, China) and used as received. Kanamycin, 6-Mercapto-1-hexanol and tris- (2-carboethyl) phosphine hydrochloride (TCEP) were purchased from Aladdin Co., Ltd. (Shanghai, China). Thiol-modified DNA aptamer sequences were synthesized by Sangon Biotechnology Co., Ltd. (Shanghai, China), purified by HPLC, confirmed by HPLC profile and mass spectrometry. These were dissolved in 1× PBS buffer (10 mM PB, 137 mM NaCl, and 2.7 mM KCl, pH 7.0) to a final concentration of 100 μM, aliquoted and stored at -20°C prior to use.

The DNA sequences used in this study were as follows.

Kanamycin-binding aptamer:

5'-HO-$(CH_2)_6$-S-S-$(CH_2)_6$-GGGACTTGGTTTAGGTAATGAGTCCC-methylene blue (MB)-3'

Doxorubicin-binding aptamer:

5'-HO-$(CH_2)_6$-S-S-$(CH_2)_6$-ACCATCTGTGTAAGGGGTAAGGGGTGGT-MB-3'

Cocaine-binding aptamer:

5'-HO-$(CH_2)_6$-S-S-$(CH_2)_6$-AGACAAGGAAAATCCTTCAATGAAGTGGGTCG-MB-3'

### Gold electrode preparation and cleaning

We fabricated gold wire electrodes using the following procedure. First, we insulated 8 cm of 0.2 mm diameter gold wire using a heat-shrinkable Teflon tubing leaving about 3 mm on one end free for DNA modification and ~1 cm at the other end for connection to the potentiostat. We then electrochemically roughened the sensor end to increase its microscopic surface area.[5] Briefly, the sensors were immersed in 0.5 M $H_2SO_4$ and rapidly pulsed between $E_{initial}$ = 0.0 V to $E_{high}$ = 2.0 V (all potentials reported in this paper are relative to Ag/AgCl) for 100 times with each pulse being of 0.02 s duration.

The electrodes were cleaned electrochemically by cycling 10 times between 0 and -1.5 V at 0.1 V $s^{-1}$ in aqueous 0.5 M NaOH using a three-electrode setup. Then the working electrodes were rinsed thoroughly with ultrapure water, transferred into 0.5 M $H_2SO_4$ and applied a chronoamperometry procedure with $E_{initial}$ = 0.0 V and $E_{high}$ = 2.0 V for 320 steps and each pulse being of 0.02 s duration. Finally, the working electrodes were transferred into 0.05 M $H_2SO_4$ using cyclic voltammetry at 0.1 V $s^{-1}$ between 0 and 1.65 V to observe integrating the area under the curve of the gold oxide reduction peak, which then dividing it by 422 μC $cm^{-2}$ to determine the electroactive area. Freshly cleaned electrodes typically exhibit a microscopic surface area of ~0.03±0.01$cm^2$.

**The fabrication of unprotected and gel-protected sensors**

The freshly cleaned electrodes were immersed in 1 μM DNA solution for 1 h at room temperature, which was previously prepared by incubating a solution of 100 μM thiolated DNA and 20 mM tris-(2-carboxyethyl) phosphine hydrochloride (TCEP) for 1 h at room temperature, and further diluted by 1×PBS buffer. Washing the sensors with deionized water and incubating in 20 mM solution containing 6-mercaptohexanol overnight at 4°C. The resulting sensors were then rinsed with deionized water, and in the case of "unprotected sensors," used as is.

Gel-protected sensors were fabricated using sensors freshly-prepared as above and then dip coated in agarose gel (Supplementary Fig. 1). To do so we first prepared the agarose-coating solutions by dissolving agarose powder (from 0.5 to 3 wt%) in 1× PBS at 95$^{o}$C, cooled the solution to 60$^{o}$C, and incubated them at this temperature for 30 min. We then gently dipped the sensors into the agarose solution 3 to 5 times with each immersion lasting ~3 s. A gel layer formed on the sensor each step, additional layers with each dip. We stored the resulting gel-protected sensors in PBS buffer prior to use.

**Diffusion model**

The diffusivity of the proteins through a cross-linked hydrogel decreases as the volume fraction of water decreases, as the size of the solute increases, and as cross-linking density increases.[1,2] In Amsden's model of this, which assumes that the diffusion of solutes is determined by the probability of solute to find an enough space between polymer fibers while taking into account the stiffness of the polymer fibers in the polymer gel,[3,4] the reduction in solute diffusivity is given by:

$$\frac{D_g}{D_0} = \exp\left[-\pi\left(\frac{r_s + r_f}{k_s\varphi^{-0.5} + 2r_f}\right)^2\right] \quad (1)$$

Here $D_0$ is the diffusion coefficient of the proteins in $H_2O$, $r_f$ represents the radius of the agarose gel fibers (1.9 nm), $r_s$ represents the hydrodynamic radius of the proteins, and $k_s$ is a scaling parameter of the system, which is constant for a given type and density of gel. φ is the volume fraction of agarose fiber in the gel, which we calculated as follows:

$$\varphi = \frac{C_{gel}}{\rho_{gel}\omega_{gel}} \quad (2)$$

Here, $C_{gel}$ is the weight fraction of agarose in the hydrogel, $r_{gel}$ is the density of dry agarose powder (1.64 g/ $cm^3$) and $w_{gel}$ is the mass fraction of agarose in the gel fiber (0.625).

$$t \approx \frac{d^2}{2D_g} \quad (3)$$

We applied equation 3 was applied to estimate the time required for diffusion of proteins within agarose gel over a given distance.

### Gel electrophoresis

We performed gel electrophoresis with whole blood and blood serum to reveal the size dependence of protein molecular diffusion in agarose gel layer. 3 wt% of agarose gel in casting tray was prepared and covered by 1x TAE buffer, the samples were loaded into the wells and the gel was electrophoresis performed for 2 h at 100 V. After electrophoresis, the gel was stained by Coomassie Blue and photographed on Tanon 5200 Chemiluminescent Imaging System (Supplementary Fig. 2).

### Confocal microscopy studies

We employed confocal microscopy to demonstrate the size dependence of molecular diffusion through a gel layer. To do so we employed the proteins BSA and fibrinogen, with the former modified with Fluorescein (emission peak at 520 nm) and the latter with Alexa Fluor 647 (emission peak at 668 nm). We fabricated a 400 μm this, 3 wt% gel slice using the protocol described above and then placed a 20 μL solution of the two proteins on top of this. We imaged protein diffusion into the slice using confocal microscopy on a Zeiss LSM 880 microscope after 1 h and 24 h. The antifouling performances of gel-protected E-AB sensors were also evaluated using fluorescein-labeled BSA and imaging with this same microscope (Supplementary Fig. 4).

**Optical microscopy studies**

We employed optical microscopy to observe the non-specific adsorption of gel-protected and unprotected sensors (Supplementary Fig. 10). We first incubated both gel-protected and unprotected sensors in whole blood for 10 h. Following this, we pealed the gel-layer off the gel-protected electrode surface and rinsed it with PBS buffer. The unprotected electrode was similarly rinsed with PBS buffer. Both were then imaged using a Zeiss Axio Scope A1.

**Electrochemical measurements**

All electrochemical measurements were performed using a multichannel CHI1040C potentiostat. In vitro experiments were performed at room temperature using a standard three-electrode cell containing a platinum counter electrode, an Ag/AgCl (3 M KCl) reference electrode and a gold wire working electrode. The in-vivo measurements were performed using a two-electrode setup in which the reference and counter electrodes were a silver wire coated with a silver chloride film.[6] All the sensors were interrogated using square wave voltammetry using a potential window of -0.05 to -0.35 V, an amplitude of 50 mV, and a frequency of 500 Hz unless otherwise noted.

**Target-binding equilibration measurements**

To determine the equilibration time constant for sensors we first interrogated them in whole blood lacking target to determine baseline peak currents. We then rapidly spiked the samples with 10 mM target via manual mixing and recorded voltammograms every ~50 s. We fit the resulting response curves to single exponential kinetics to determine the equilibration time constants.

**Target titration measurements**

We performed target titrations to investigate whether the gel coating has an impact on the dynamic range of gel-protected sensors in comparison to that of unprotected ones. To do so we first interrogated both unprotected and gel-protected sensors in target-free blood sample to determine baseline peak currents. Following this, we spiked the blood with monotonically increasing target concentrations, and recorded voltammograms 1 min after each addition of target.

**In vivo experiments**

**Animals**. 8-week old Sprague-Dawley (SD) rats and 4-week old BALB/C nude mice were purchased from Beijing Vital River Laboratory Animal Technology Co., Ltd. All animals were acclimatized for 1 week at 26°C, 40% humidity and 12 h light dark cycle room, and sufficient water and food were ensured. The animal ethics involved in the study were approved by Tongji Hospital, Tongji Medical College, Huazhong University of science and technology.

**Anesthesia**. Before the sensor implantation, we intraperitoneally injected the animals with 5% chloral hydrate (1 mL per 100 g body weight), with simultaneous monitoring of the heart rate and respiration of rats and mice to ensure that appropriate depth of anesthesia was achieved.

**Venous sensor implantation**. Prior to the implantation of intravenous sensors in the jugular, we first injected heparin (1000 U/mL, 0.2 mL) into the left jugular vein to reduce clotting. Then we performed a detachable ligation at the distal end of the right jugular vein, and cut a small incision at the proximal end. Following on this, we implanted the sensors into the right jugular vein along the small opening of this cut, and tired the vessel wall together with the sensor to secure it in place. We then used the left jugular vein for injection of the target and for blood sample collection.

**Bladder sensor implantation**. We first opened the abdominal cavity at the midline of the lower abdomen, with the bladder exposed for sensor implantations. We then inserted a 24G indwelling needle into the bladder, with the indwelling part of this needle placed in the bladder. Following on this, we insert the sensor into the bladder through the external opening of the indwelling needle, while allowing urine to flow out through the indwelling needle.

**Muscle sensor implantation**. For this we employed an implantation device composed of two parts: an outer, elastic catheter tube and an inner steel needle. We first inserted both into the muscle, and then removed the inner steel needle, leaving the outer catheter inside of the muscle to provide an access channel. We inserted the sensor through the catheter and then removed the catheter, leaving the sensor implanted in the muscle.

**Tumor sensor implantation**. We first transplanted $10^6$ HeLa cells into the right armpit of 5-week-old BALB/C nude mice. After the tumor size reached to about 200 $mm^3$, we then implanted sensors into the tumor via the procedure described above for muscle implantation.

**HPLC-ELSD measurements**

**Chromatography.** We prepared samples by adding 150 μL trichloroacetic acid and 850 μL of ultrapure water was added to a 250 μL blood sample. After addition, the sample was vortexed for 1 min and then centrifuged 10 min at 12,000 rpm. The supernatant was collected and passed through a 0.45 μm filter. We injected 20 uL of the resulting solution into HPLC (Beijing QingBoHua HPLC, Beijing QingBoHua Technologies co., ltd. China) modular system with an ELSD (Evaporative Light Scattering Detection, UM-4800, unimicro (shanghai) technologies co., ltd.) detector. The column temperature was set to 25°C, the air pressure in the ELSD detection module was set to 3.5 bar, and the evaporation temperature to 40°C. The mobile phase consisted of aqueous phase (with 0.3% TFA) and methanol (5:95, v/v) at a flow rate of 1 mL/min. Each measurement was repeated at least three times to determine confidence intervals.

We constructed an HPLC calibration curve by plotting observed peak areas against the known drug concentrations of standard solutions prepared at 10, 20, 50, 100, 200, 500, 800, and 1000 μg/mL by diluting the stock solution (100 mg/mL in PBS buffer) with calf blood. Typical chromatogram of kanamycin sample from ELSD is shown in Supplementary Fig. 14 (3.65 min). After logarithmic transformation, the data provide a linear calibration following the equation: $\log A = 1.944 + 0.218 \times \log C$, where $A$ is the value of the peak area, $C$ is the value of kanamycin concentration. The correlation coefficient ($r^2 \geqslant 0.97$) value indicated a suitable correlation between the concentrations of kanamycin and the peak area.

**Biocompatibility assessment**

**Implantations**. We randomly divided 25 eight-week old SD rats into 5 groups: control group, tail vein (no gel) group, tail vein (gel) group, muscle (no gel) group and muscle (gel) group. We performed all the surgical procedures for the control group, without any sensor implantations. We implanted E-AB sensors into tail vein or muscle in other four groups, leaving the sensors in the body for one week. During this week, we recorded the body weight, water/food consumption and behavior of all animals. At the end of one week, we collected the blood, tissues (tail vein and muscle), heart, liver, spleen, lung and kidney from all five groups for blood examinations and histopathological studies.

**Histopathological examination**. First, we observed the freshly-obtained tissues or organs by naked eyes, observing no thrombosis in the blood vessels or other organs. Then, we performed hematoxylin&eosin staining on the samples and imaged them to determine their morphological and pathological characteristics.

**Blood test**. All blood samples were stored in anticoagulant tubes for 4°C, and blood cells were classified within 24 hours.

**ELISA assays**. All blood samples were centrifuged at 10,000 rpm (4°C, 10 min) to obtain plasma for subsequent detection of inflammatory factors and thrombus factors. The inflammatory factors and thrombus factors detected included CRP (70-EK394-96, MultiSciences, China), IL-12 (EK1652, BOSTER, China), IL-6 (70-EK306/3-96, MultiSciences, China), TNF-α (70-EK382/3-96, MultiSciences, China), ET (EK0952, BOSTER, China), TM (E-EL-R0960c, Elabscience, China) and D-Dimer (E-EL-R0317c, Elabscience, China). All ELISA assay were performed in accordance with the manufacturer's instructions.

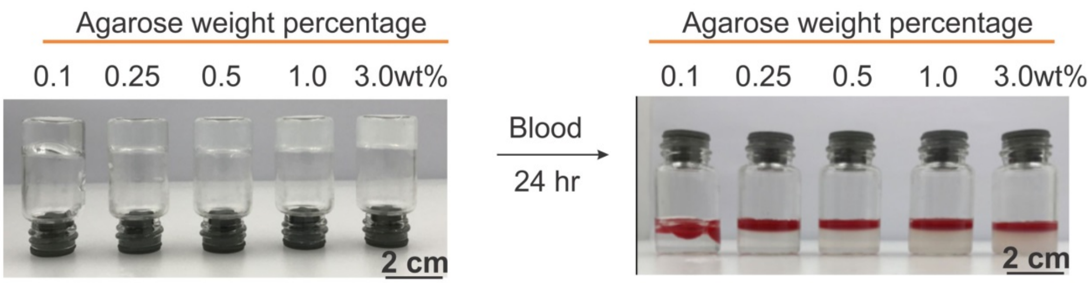


**Supplementary Fig. 1** Optimization and characterization of agarose gel. We prepared a series of agarose gel solution in the range from 0.1 wt% to 3 wt%. (left) All form stable gels except 0.1 wt%. (right) Not surprisingly higher (≥ 0.5 wt%) concentration gels better resisted the penetration of blood cells.

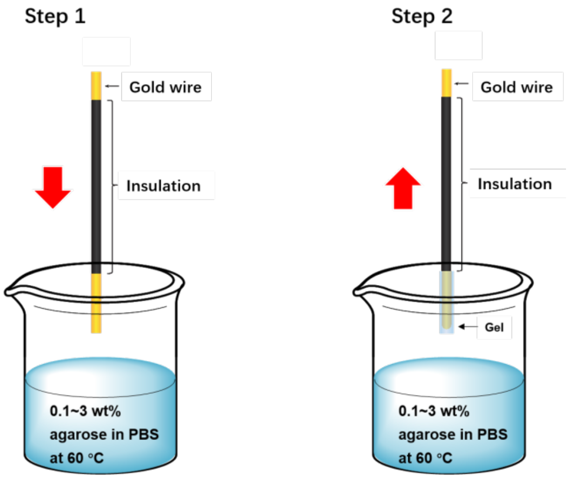


**Supplementary Fig. 2** We prepared the gel-protected sensors via a dip-coating procedure. (1) We firstly prepared the agarose-coating solution by dissolving agarose powder in 1× PBS at 95°C; (2) Then we stored the agarose solution in an incubator at 60°C until a highly viscose coating solution was obtained; (3) We gently and slowly dipped the sensors into the coating solution 3 to 5 times, immersing them for ~3 s each time.

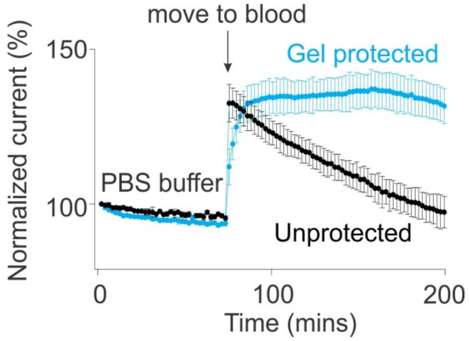


**Supplementary Fig. 3** While the gel-protected and unprotected sensors exhibit a similar stability in simple buffers, their stabilities differ dramatically when these two are placed in whole blood. While unprotected sensors lose up to 40% of their signal over ~100 min in whole blood, gel-protected sensors exhibit less than 5% signal loss under these same conditions.

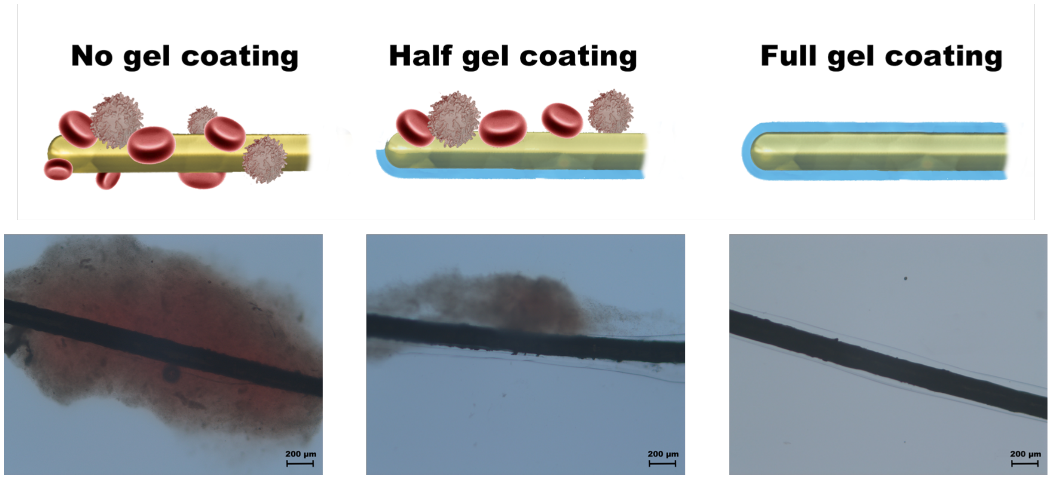


**Supplementary Fig. 4** Optical microscopy study demonstrated that the gel-coating protected E-AB sensors from fouling. Specifically, (left) after 10 h incubation in whole blood, we observed a large blood clot absorbed onto an unprotected sensor. (middle) When we removed half of the gel coating from a sensor, we observed blood adsorption on the unprotected side and no adsorption on the other. (right) A gel protected sensor does not exhibit any significant adsorption.

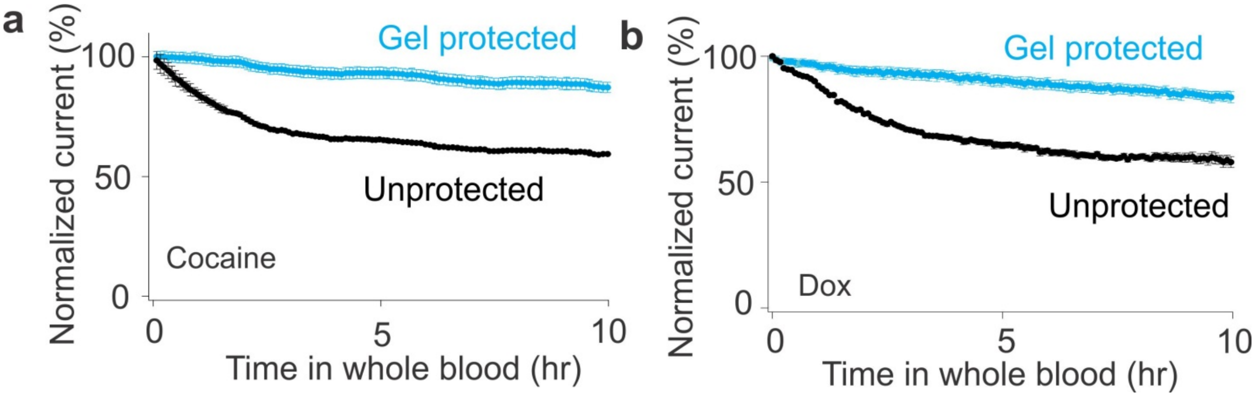


**Supplementary Fig. 5** In addition to improving the stability of sensors against kanamycin (Fig. 3d, the improved stability associated with gel-protected sensors is also seen for sensors employing aptamers against (A) cocaine and (B) doxorubicin. Once again, while the unprotected sensors exhibited a significant signal decrease over 10 h in whole blood, the gel-protected sensors exhibit less than 10% signal loss under these same conditions.

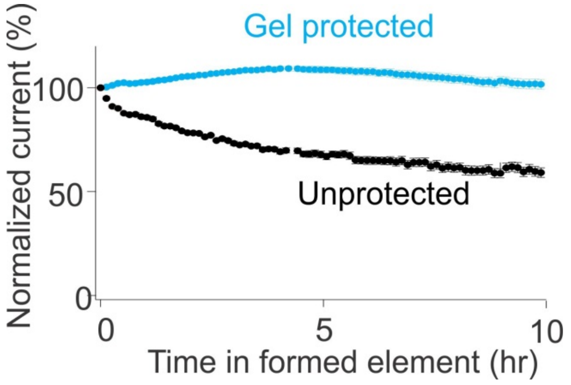


**Supplementary Fig. 6** When interrogated in formed element samples over the course of 10 h, unprotected kanamycin sensors exhibit 50% signal loss. Gel-protected sensors, in contrast, exhibit less than 5% signal loss.

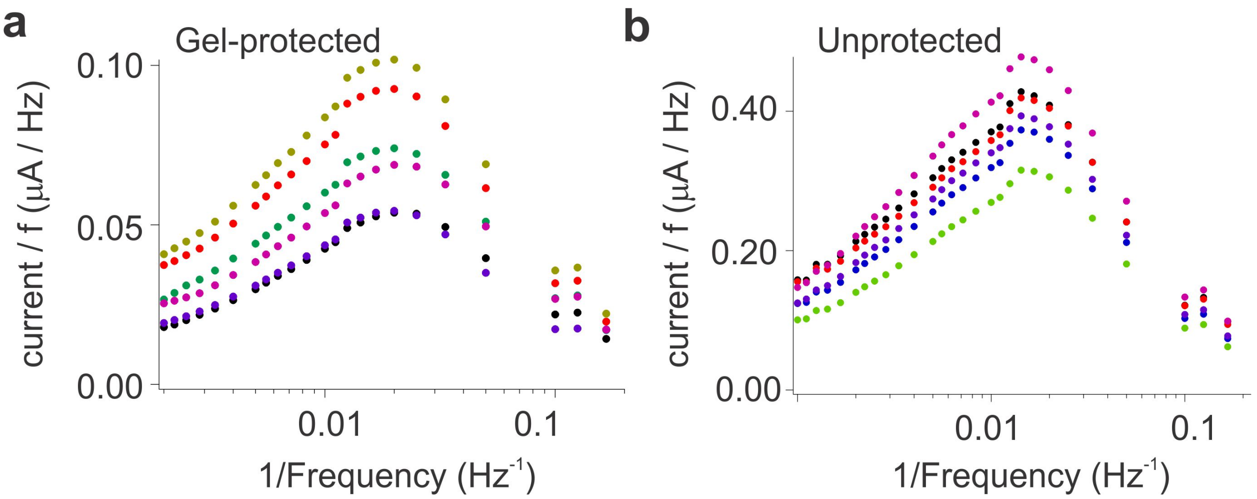


**Supplementary Fig. 7** Target binding in E-AB sensors is coupled to a change in the conformation and flexibility of the aptamer, which, in turn, changes the efficiency with which electrons are transferred to or from the electrode. As observed here (via Laver on plots [7]), the difference in electron transfer rates between gel-protected and unprotected sensors is negligible. Each panel includes data collected using six individual, hand-fabricated sensors to illustrate the good reproducibility of the electron transfer kinetics. The variation in peak height simply reflects variability in the total surface area of each sensor and, with that, variability in the total charge transfer,

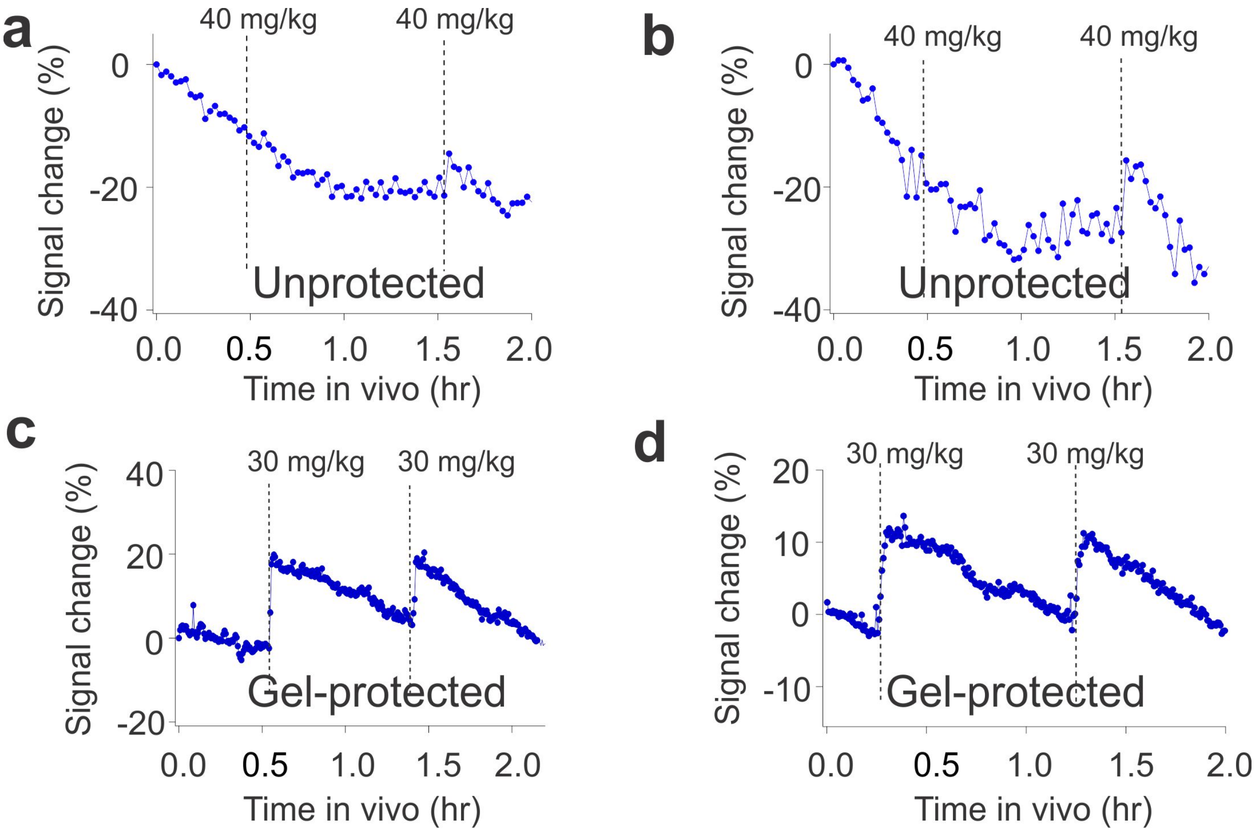


**Supplementary Fig. 8** Gel-protected E-AB sensors support continuous, real-time molecular measurements in situ in the living body without the use of the drift-correction algorithms previously used in such deployments. To show this, we first emplaced kanamycin-detecting sensors in the external jugular veins of anesthetized Sprague-Dawley rats, which we then challenged via intravenous injection of the drug into the opposite external jugular vein. (a) and (b) Unprotected sensors exhibited significant drift over the course of these few-hour experiments, rendering them incapable of determining the target concentrations without employing drift correction methods. (c) and (d) In contrast, gel-protected sensors are much more stable, achieving precise, continuous molecular measurement. These data are replicate measurements in the manuscript (Fig. 4), illustrating the reproducibility of E-AB sensors for in-vivo measurements.

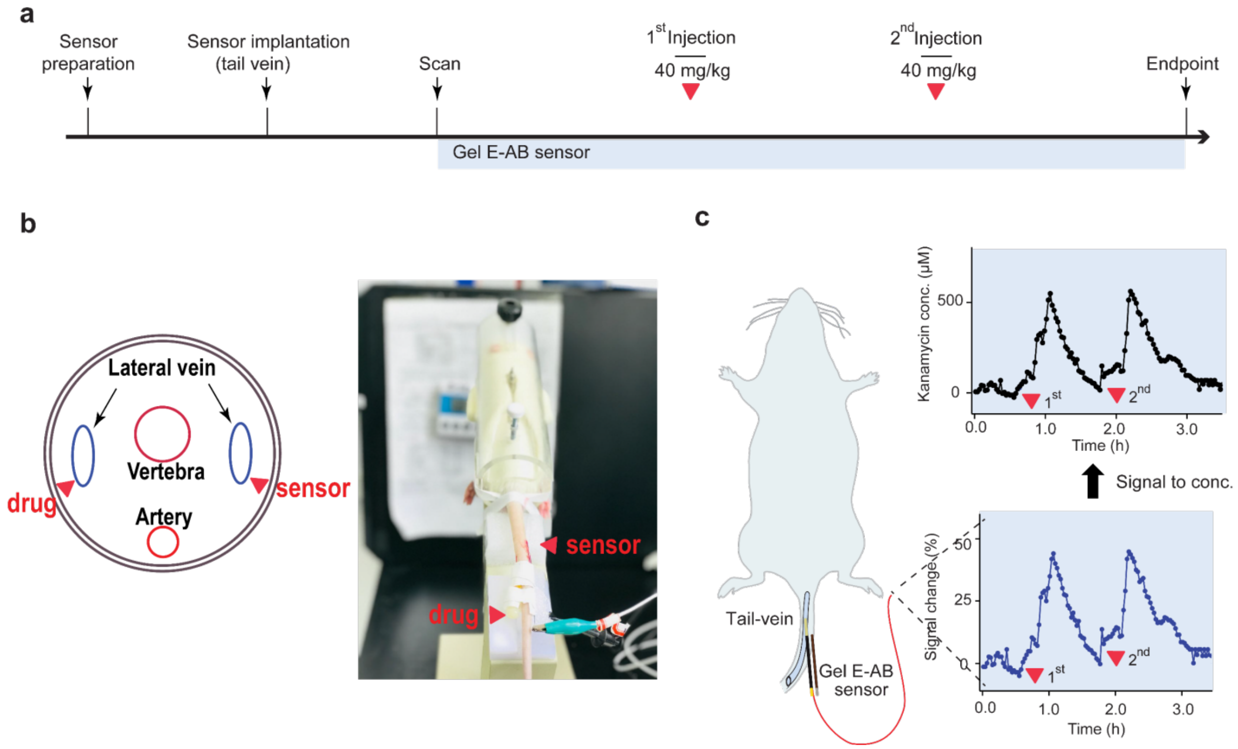


**Supplementary Fig. 9** In the initial assessment, we implanted our gel-protected sensors in the tail veins of live rats. (a) We followed the standard protocol for sensor implantations and E-AB sensor measurements. (b) The illustrations of E-AB implantations and drug infusion. We implanted E-AB sensor on one of lateral vein of the tail, and infused the drug on the other of lateral vein. (c) Upon injecting two, sequential 40 mg/kg doses (a human therapeutic level), we observed two clear, corresponding peaks in drug concentration.

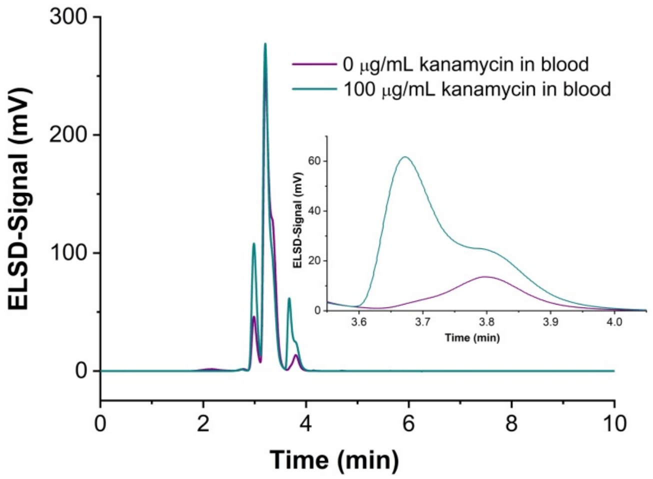


**Supplementary Fig. 10** Typical chromatogram used for the determination of kanamycin in rat blood.

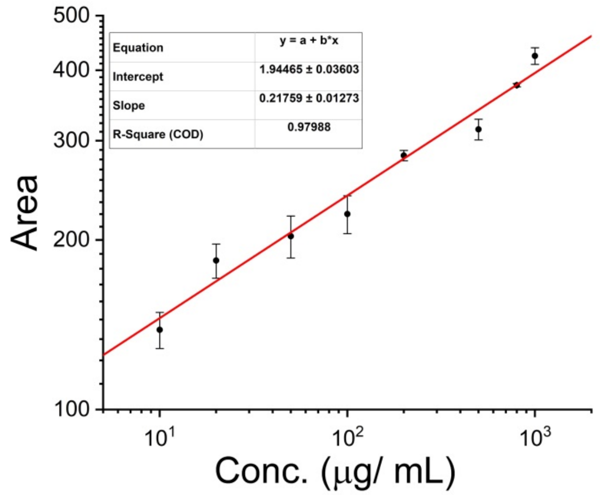


**Supplementary Fig. 11** The HPLC-ELSD calibration curve used for the ex-vivo quantification of kanamycin in blood.

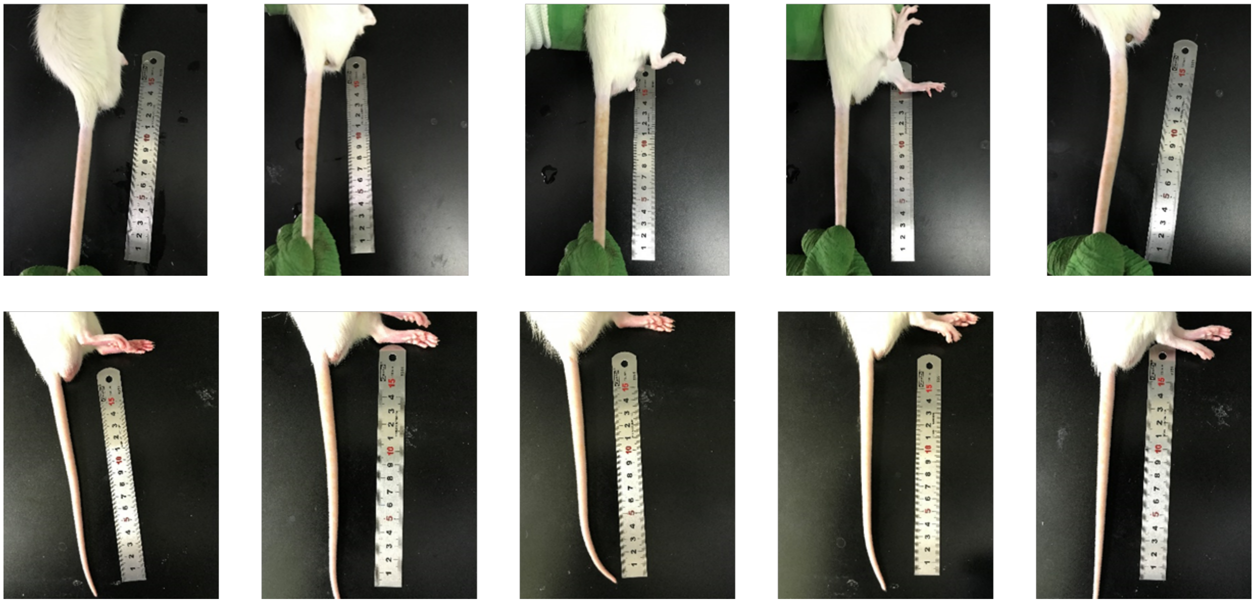

**Supplementary Fig. 12** From the photographs of the tail vein in control rat group at day 1 (top) and day 7 (bottom), we observed complete recovery from the implantation surgery and no inflammation of the skin.

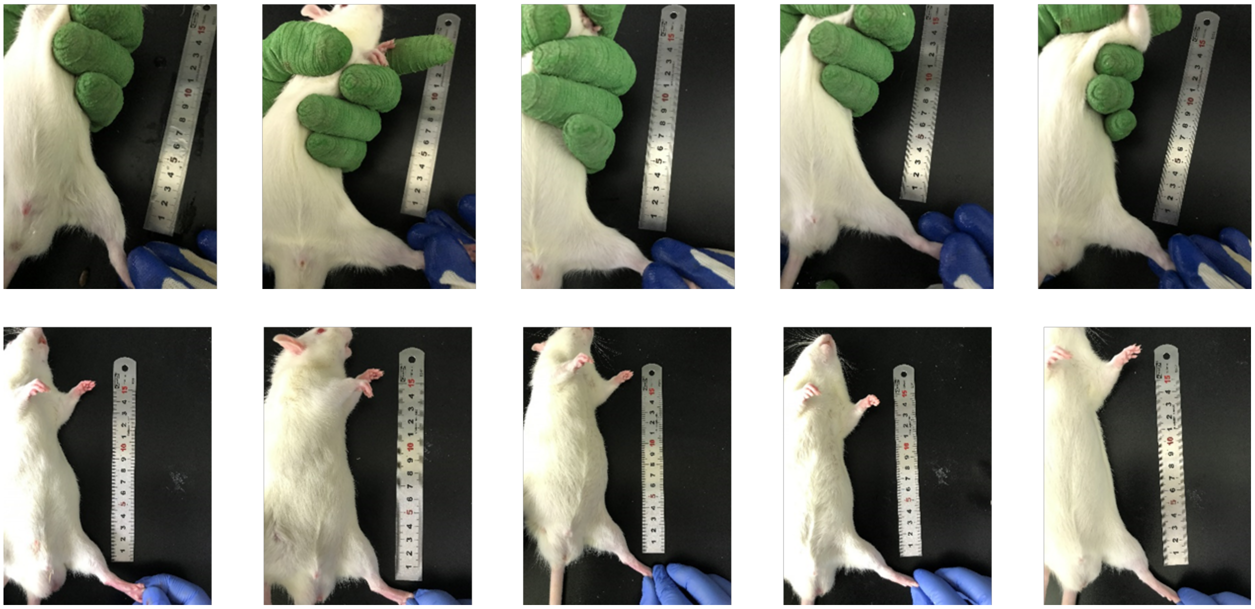

**Supplementary Fig. 13** From the photographs left hind limb in control rat group at day 1 (top) and day 7 (bottom), we observed complete recovery from the implantation surgery and no inflammation of the skin.

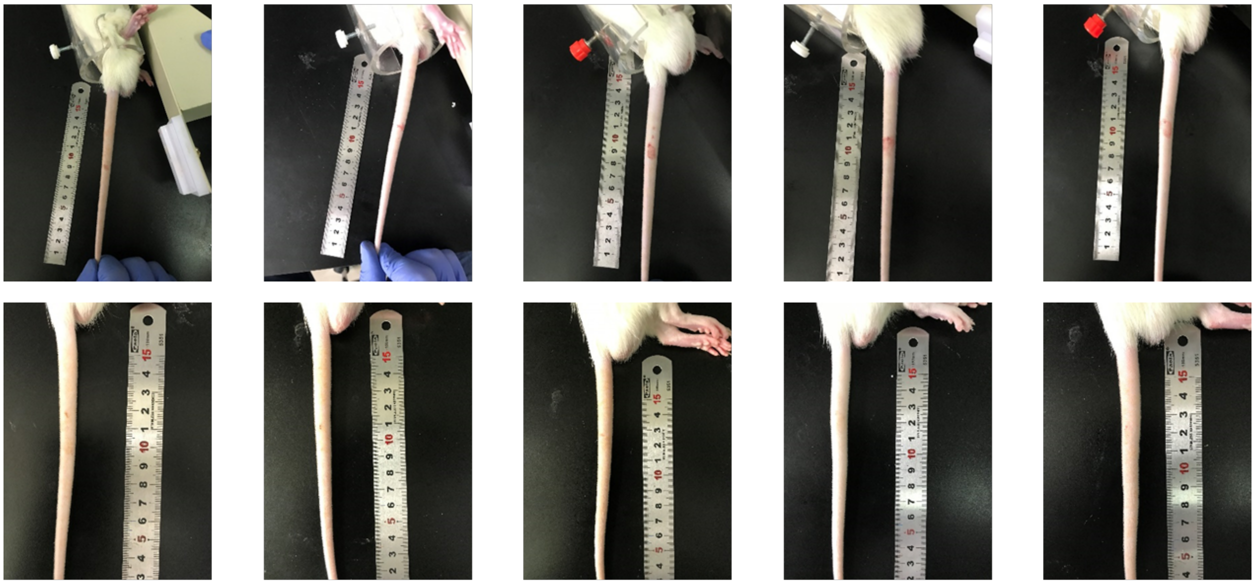

**Supplementary Fig. 14** From the photographs of the tail vein implanted with unprotected sensors at day 1 (top) and day 7 (bottom), we observed complete recovery from the implantation surgery and no inflammation of the skin.

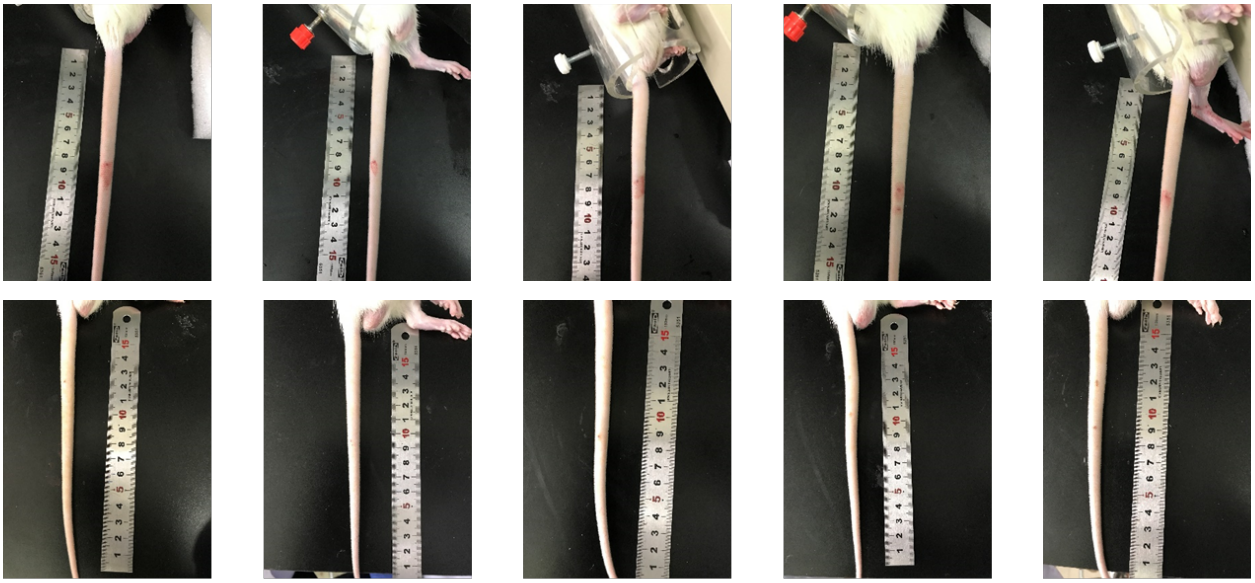

**Supplementary Fig. 15** From the photographs of the tail vein implanted with gel-protected sensors at day 1 (top) and day 7 (bottom), we observed complete recovery from the implantation surgery and no inflammation of the skin.

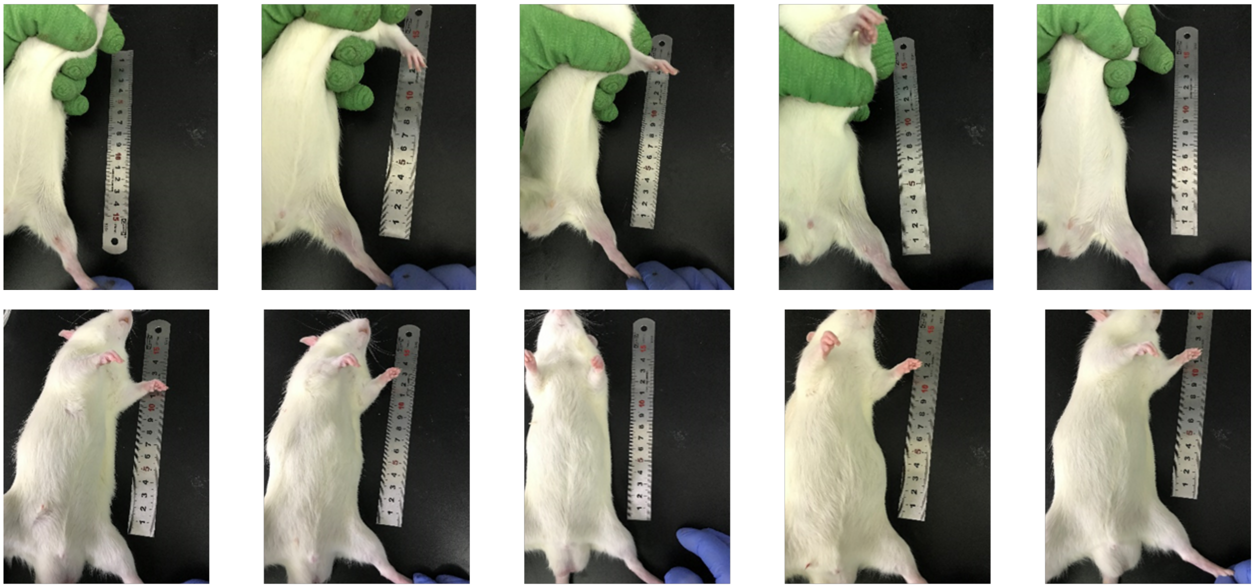

**Supplementary Fig. 16** From the photographs of the left hind limb implanted with unprotected sensors at day 1 (top) and day 7 (bottom), we observed complete recovery from the implantation surgery and no inflammation of the skin.

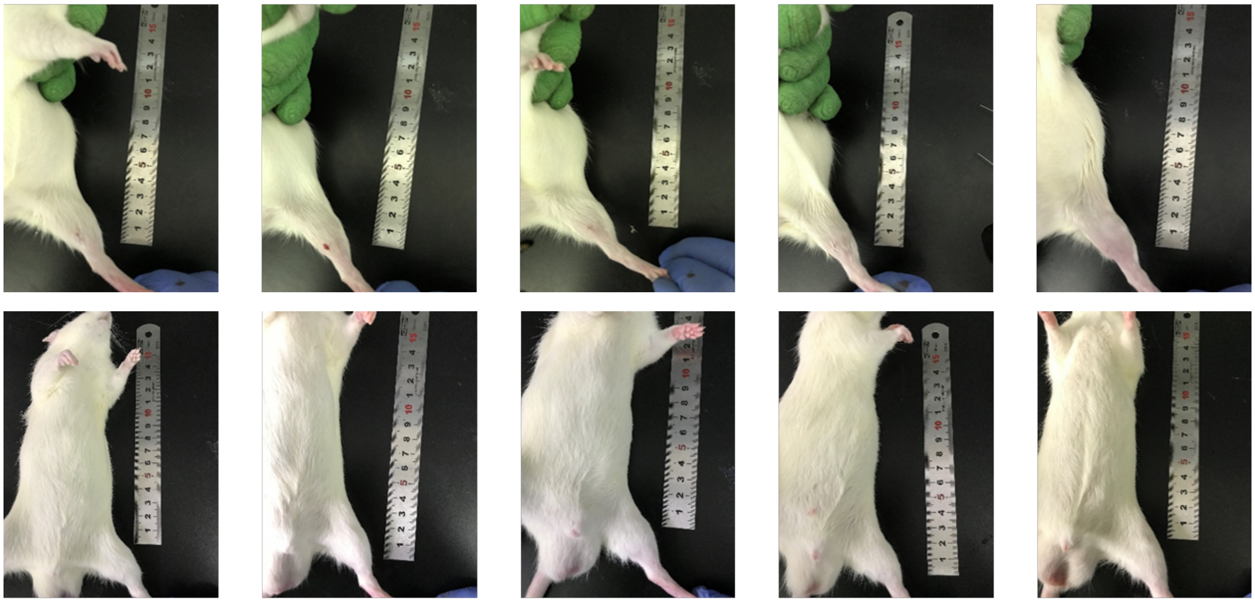

**Supplementary Fig. 17** From the photographs of the left hind limb implanted with gel-protected sensors at day 1 (top) and day 7 (bottom), we observed complete recovery from the implantation surgery and no inflammation of the skin.

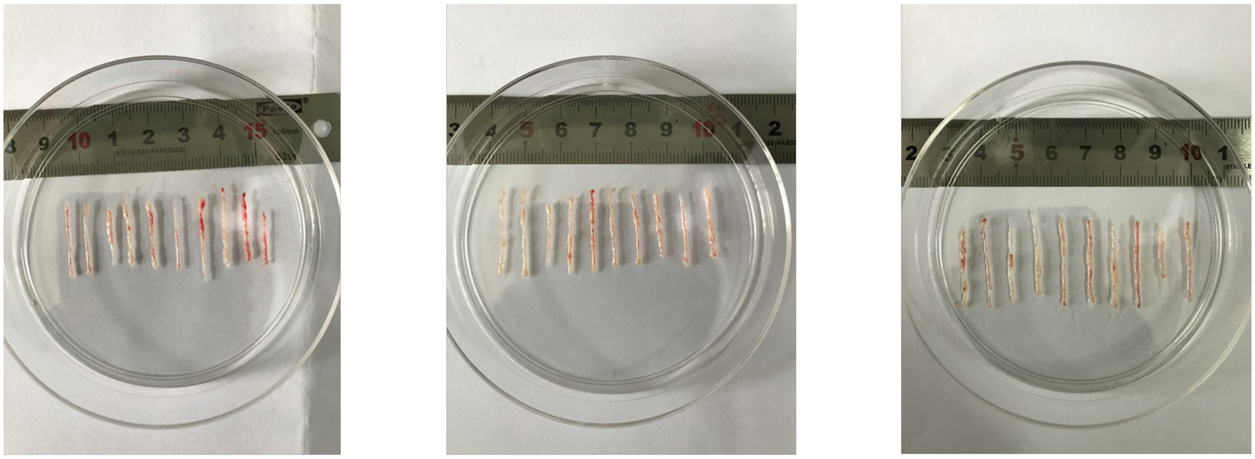

Tail vein (control) Tail vein (no gel) Tail vein (with gel)

**Supplementary Fig. 18** The photograph of tail vein from three groups of rats at day 7: (left) control group without implantation; (middle) implantation group with unprotected sensors; (right) implantation group with gel-protected sensors. No significant thrombosis was observed in any of the three groups.
